# Validity and internal consistency of EQ-5D-3L quality of life tool among pre-dialysis patients with chronic kidney disease in Sri Lanka, a lower middle-income country

**DOI:** 10.1101/524603

**Authors:** Sameera Senanayake, PKB Mahesh, Nalika Gunawardena, Nicholas Graves, Sanjeewa Kularatna

## Abstract

**Objective:** EQ-5D-3L is a generic QOL tool used mainly in economic evaluations. Burden of Chronic Kidney Disease (CKD) is rising in Sri Lanka. Assessing the validity of generic QOL tools creates new opportunities of their utilization among patients with CKD.

**Methods:** A cross-sectional study was conducted among 1036 CKD patients, selected using the simple random sampling technique. The validity was tested with six a-priori hypotheses. These included construct validity assessments, evaluating convergent validity and performing known group comparisons. EQ-5D-3L, Short Form-36 (SF-36), Center for Epidemiological Studies Depression Scale (CES-D-20) and General Health Questionnaire-12 (GHQ-12) were used to assess QOL, presence of depression and psychological distress respectively. Internal consistency of the whole tool and when each item is removed was assessed by Cronbach alpha.

**Results:** The response rate was 99.2%. Majority of participants were males (n=646,62.4%) in the age category of 41-60 (n=530; 51.2%). Most were in either stage 4 or 5 of CKD (n=646,75.1%). The summary measures of SF-36, positively and significantly correlated with the EQ-5D-3L index and VAS scores (p<0.001). EQ-5D-3L QOL scores were significantly different between the group with depression and without as measured by CES-D-20 (p<0.001). Assessed using GHQ-12, similar significance was detected between the group with psychological distress and without (p<0.001). The Cronbach alpha was 0.834 and when each item was removed, ranged from 0.782 to 0.832.

**Conclusion:** EQ-5D-3L is a valid generic QOL tool with satisfactory internal consistency to be used among CKD patients in the pre-dialysis stage.

## Introduction

Chronic Kidney Disease (CKD) has become a major global burden of disease accounting for a significant mortality and disability adjusted life years (1). In the Global Burden of Disease Study-2015, it was ranked as the “12^th^ most common cause of death” with an increased overall mortality over 30% in the previous decade (2). This rise has been “alarming” in the low- and middle-income settings (3). Additional emphasis on CKD has been recommended in relation to these settings owing to the existing deficiencies in health policies and practices (2).

Sri Lanka which is an agricultural lower-middle-income country, is affected by the rising trend of chronic non-communicable diseases (NCDs) due to demographic and epidemiological transitions (4,5). CKD has become a major burden of the healthcare system in Sri Lanka (6). With the contribution of occurrence of unexpected types of it like CKD of unknown etiology as well as due to the concomitant increase of incidence of other NCDs, its burden is expected to rise further (6,7). In addition, CKD impose numerous social and economic threats in Sri Lanka(8).

Quality of life (QOL) reflects a person’s subjective evaluation of his or her position of life in the living contexts (4). The QOL related to impact of health conditions is referred to as health-related QOL (9). Measurement of QOL become utmost important among patients with CKD, due to the problems they are forced to face by the disease condition. They face a diverse range of symptoms, distresses and even depression (1,10). Furthermore, the condition would make them suffer from social and financial inabilities as well. They need to be in a potential waiting period for getting the dialysis done, even when they enter that stage.

Health-related QOL can be measured by using generic as well as disease specific tools (11). SF-36 and EQ-5D-3L are such generic QOL tools (12). Assessment of validity of QOL tools, for specific disease conditions would create many opportunities for utilization within specific settings (13). This is particularly beneficial for generic QOL tools, as this allows QOL comparisons between different conditions (14). Validity measures to what extent the tool measures what it is expected to measure (15,16). Reliability reflects the reproducibility of the same concept when repeated measures are observed. It can be assessed by methods including internal consistency and test-retest method (17).

In the absence of gold standard tests for measurements like QOL, the validity assessments could be done for construct validity with methods such as “convergent/divergent validity assessments” and “known group comparisons” (13,15,18). The EQ-5D-3L health states have been valued using preference of general population in Sri Lanka(19). Furthermore using it, health related QOL has been assessed in relation to several main chronic NCDs in Sri Lanka (4, 20). Yet the validity of it, in relation to CKD has not been assessed.

The purpose of this study is to assess the validity and the internal consistency of EQ-5D-3L among patients with kidney diseases who are in their pre-dialysis stage.

## Materials and Methods

### Selection of the sample

A descriptive cross-sectional study was conducted in Anuradhapura district of Sri Lanka. This district, located in the North Central Province, is experiencing a rising trend of CKD. Patients with CKD confirmed by a medical specialist based on the recommended guidelines (ie. Glomerular Filtration Rate (GFR) being less than 60ml/min per 1.73m^2^ body surface area in two measurements made three months apart) comprised the study population. All patients above 18 years were included. Patients who had renal transplantations were excluded.

The sample size (n) for the estimation of the construct validity was based on the publication of Cohen (1992) for the “product-moment correlation coefficient”(21). Cohen (1992) recommends a sample size of 783 for a small effect size (21). With an assumed response rate of 75%, minimal sample size was calculated as 1044. Patients with CKD in the district of Anuradhapura are registered in a population-based CKD register according to the geographic health units in which they reside. The district comprise 19 health units and the required number of study units were allocated to the 19 health units in proportionate to the number of registered patients in each health unit. Using the population based register as the frame, simple random sampling technique was used to select the study units from each the 19 health units within the district.

### Assessment of validity and internal consistency

The construct validity of the EQ-5D-3L was assessed using the following six a-priori hypotheses.

1. The EQ-5D-3L index scores would significantly correlate with SF-36 summary scores in the positive direction with acceptable strength
2. The EQ-5D-3L VAS scores would significantly correlate with SF-36 summary scores in the positive direction with acceptable strength
3. The EQ-5D-3L index scores would be significantly different between the groups with depression and without depression
4. The EQ-5D-3L VAS scores would be significantly different between the groups with and without depression
5. The EQ-5D-3L index scores would be significantly different between the groups with depression and without depression
6. The EQ-5D-3L VAS scores would be significantly different between the groups with and without psychological distress

The internal consistency was assessed using Cronbach alpha measurement for all the items as well as when each item is removed.

EQ-5D-3L developed by the EuroQol group consist of a descriptive system and a Visual Analogue Scale (VAS). The descriptive system comprises of five questions (on mobility, self-care, daily activities, pain and mood) with each carrying three levels of possible responses. Each level reflects a level of impairment as “with no problem”, “with some problems” and “with severe problems”. An index score can be obtained from these questions which is expressed as a negative or positive fraction. VAS may range from 0 to 100 and reflects the general health as perceived by the participant (22). EQ-5D-3L has been previously translated to Sinhalese language and population norms has been developed in 2014 (19). Other tools used comprised interviewer administered forms of SF-36, CES-D-20 and GHQ-12, in addition to the tool to assess socio-demographic and medical details.

SF-36 was used in the assessment of a-priori hypothesis 1 and 2. It includes 36 items out of which 35 are used to calculate eight QOL domain scores. Four each of these domains give rise to the “physical-component” and “mental-component” summary measures. SF-36 focuses on the period of previous 28 days in getting the responses (23). Its validity has been ensured in Sri Lankan setting for several conditions (24,25). Furthermore it has been used in this setting to explore QOL of several conditions (4,20).

The presence of depression was screened by the CES-D. It has 20 questions and a maximum score of 60 is allocated. A score above 15 is indicative of depression. Its sensitivity is over 80% and the specificity is over 90% in relation to the local setting (26). In screening for psychological distress, GHQ-12 questionnaire was used. It gives a maximum score of 12. A cut-off value of two or more has been recommended to the local setting with a sensitivity and specificity more than 70% (27).

### Data analysis

Since the normality assessment showed non-normal distributions non-parametric techniques were used in the analysis. For the a-priori hypothesis 1 and 2, Spearman correlation coefficient was used(28). For the a-priori hypotheses No. 3 to 6, Mann Whitney U test was used(29). The significance level was considered as 5%, but when lower p values were observed, relevant significant levels were reported.

Ethics approval was obtained from the Ethics Review Committee of the Faculty of Medicine, University of Colombo prior to the data collection (EC-15-081). Informed written consent was taken from participants.

## Results

The response rate was 99.2%. The median (IQR) age of the participants was 59 (52 to 66) years. More than half (51.2%) the population were between 41-60 years old. The majority were males (62.4%) and were unemployed (64.7%). Nearly four fifth were in stages of 4 or 5 in the CKD staging. Of the population 64.2% were screening positive for depression while 74.4% were positive for psychological distress (Table 1).

**Table 1:** Characteristics of the participants.

| Characteristic (N=1036) | Frequency | Percentage (%) |
| --- | --- | --- |
| Age (Years) |  |  |
| 18 - 40 | 39 | 3.8 |
| 41 - 60 | 530 | 51.2 |
| 61- 80 | 467 | 45.1 |
| Gender |  |  |
| Male | 646 | 62.4 |
| Female | 390 | 37.6 |
| Highest education level |  |  |
| No formal education | 73 | 7.0 |
| Up to Grade 5 | 398 | 38.4 |
| From Grade 6 to 11 | 361 | 34.8 |
| Passed GCE (O/L) | 173 | 16.7 |
| GCE (A/L) and above | 31 | 3.0 |
| Employment status |  |  |
| Not employed | 670 | 64.7 |
| Employed | 366 | 35.3 |
| Presence of co-morbidities |  |  |
| Comorbidities present | 735 | 70.9 |
| No comorbidities | 301 | 29.1 |
| CKD stage |  |  |
| Early stage | 258 | 24.9 |
| Stage 4 | 626 | 60.4 |
| Stage 5 | 152 | 14.7 |
| Depression status |  |  |
| Depressed | 665 | 64.2 |
| Not depressed | 371 | 35.8 |
| Psychological distress status |  |  |
| Distressed | 772 | 74.5 |
| Not distressed | 264 | 25.5 |

The descriptive system of the EQ-5D-3L comprises of five questions (on mobility, self-care, daily activities, pain and mood) with each carrying three levels of possible responses. Except in the self-care domain, in all other four domains, the majority had stated level 2 or 3 responses (i.e. either having some or major problems). For the pain and the mood domains, responses with some-or-severe problems were reported by more than three fourth. In addition, in more than half of the participants, the mobility and the usual activities have been affected to a greater or lesser degree (Table 2).

**Table 2:** Distribution of responses for the descriptive question of the EQ-5D-3L.

| Item | Response |  |  |
| --- | --- | --- | --- |
|  | No problem | Some problem | Severe problem |
|  | N (%) | N (%) | N (%) |
| Mobility | 500 (48.3) | 519 (50.1) | 17 (1.6) |
| Self-care | 623 (60.1) | 388 (37.5) | 25 (2.4) |
| Usual activities | 462 (44.6) | 544 (52.5) | 30 (2.9) |
| Pain | 175 (16.9) | 714 (68.9) | 147 (14.2) |
| Mood | 253 (24.4) | 652 (62.9) | 131 (12.6) |

Role-limitation-emotional domain had recorded lowest scores out of the eight domains of the SF-36. The mental-component summary score has recorded a slightly higher score than that of the physical-component. Mean EQ-5D-3L index score was 0.52 (SD 0.33). Median scores of all the eight domains of both physical health and mental health summary components had the scores below 50.0. The highest mean score was for the Vitality domain (45.29; SD 11.53) while the lowest was for the Role-limitation-emotional (23.94; SD 33.25) (Table 3).

**Table 3:** QOL scores for domains of SF-36 and EQ-5D-3L.

| **Domain** | **Mean (SD)** | **Median (IQR)** |
| --- | --- | --- |
| SF-36 General Health | 38.17 (19.33) | 30.00 (25.00- 50.00) |
| SF-36 Physical functioning | 42.02 (23.58) | 40.00 (25.00- 60.00) |
| SF-36 Pain | 36.86 (16.58) | 42.50 (22.50- 45.00) |
| SF-36 Role-limitation-physical | 27.26 (34.12) | 0.00 (0.00- 50.00) |
| SF-36 Role-limitation-emotional | 23.94 (33.25) | 0.00 (0.00- 33.33) |
| SF-36 Vitality | 45.29 (11.53) | 50.00 (40.00- 53.75) |
| SF-36 Social functioning | 42.78 (22.03) | 50.00 (25.00- 62.50) |
| SF-38 Emotional well being | 48.38 (9.33) | 48.00 (44.00- 52.00) |
| SF-36 Physical summary score | 36.08 (14.70) | 35.00 (26.25 – 42.50) |
| SF-36 Mental summary score | 40.10 (12.10) | 39.10 (34.83 -43.09) |
| EQ-5D-3L index score | 0.52 (0.33) | 0.58 (0.34 – 0.75) |
| EQ-5D-3L VAS score | 51.35 (16.88) | 50.00 (40.00- 60.00) |

The Spearman correlation coefficients between SF-36 summary measures versus the EQ5D-3L index and VAS scores were statistically significant (p<0.001) as well as positive in direction, even though being lower in strength (Table 4). The index values showed relatively stronger associations than the VAS scores.

**Table 4:** Correlation of SF-36 summary scores with EQ-5D-3L scores.

|  | **EQ-5D-3L index score**<br>**Spearman rho (p)** | **EQ-5D-3L VAS score**<br>**Spearman rho (p)** |
| --- | --- | --- |
| SF-36 Physical summary score | 0.28 (<0.001) | 0.21 (<0.001) |
| SF-36 Mental summary score | 0.34 (<0.001) | 0.18 (<0.001) |

The comparison of the EQ-5D-3L and SF-36 scores between the depressed and non-depressed groups were carried out to test the 5^th^ a-priori hypotheses (Table 5). It shows that statistically significant differences are observed between the QOL measures between the two groups as detected by both the tools (p<0.001).

**Table 5:**
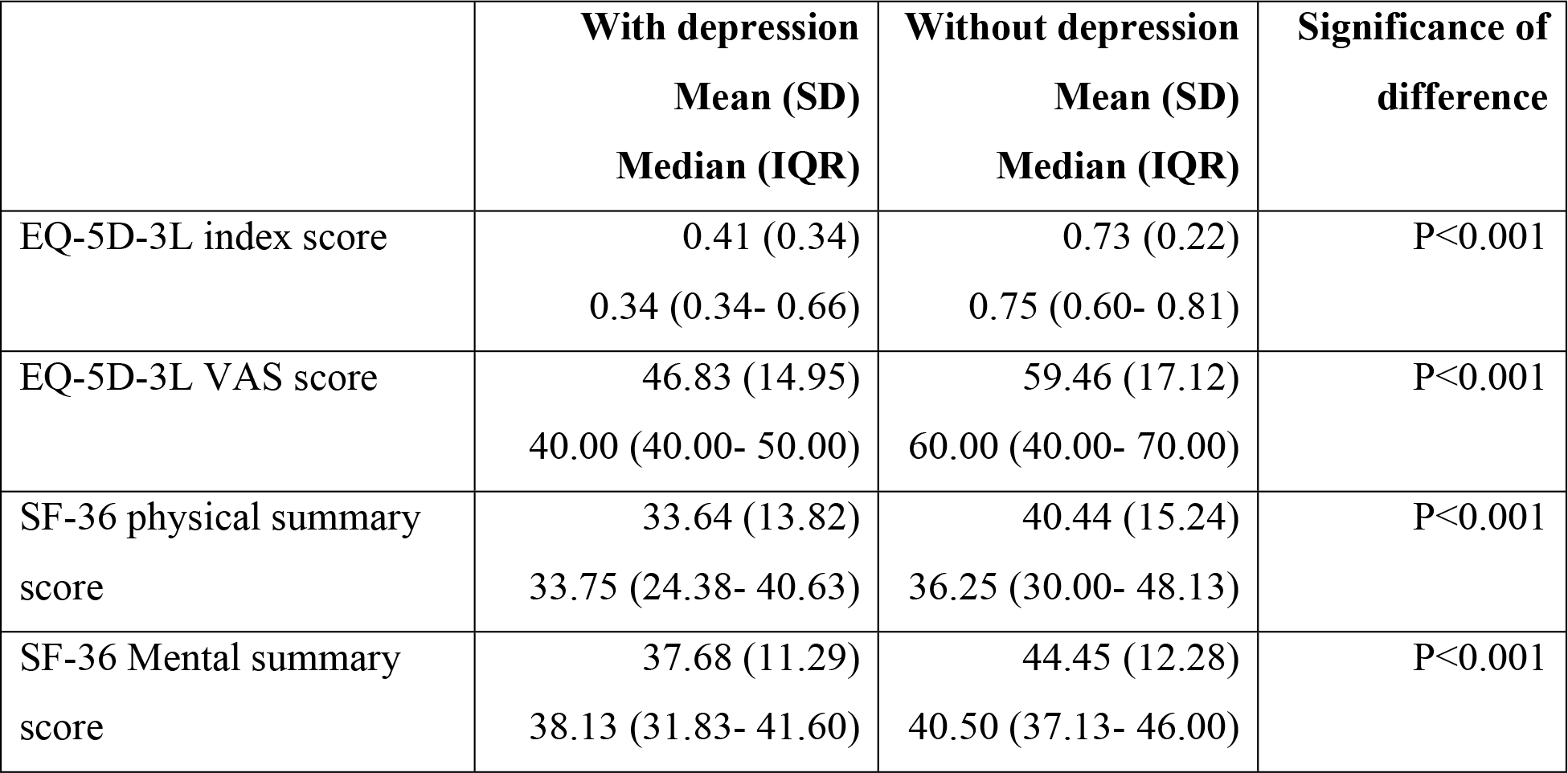
Known group comparison among participant with or without depression with SF-36 summary scores and EQ-5D-3L scores.

Similar statistically significant differences are shown between the groups with and without psychological distress (p<0.001) (Table 6).

**Table 6:** Known group comparison among participant with or without psychological distress with SF-36 summary scores and EQ-5D-3L scores.

|  | **With distress**<br>**Mean (SD)**<br>**Median (IQR)** | **Without distress**<br>**Mean (SD)**<br>**Median (IQR)** | **Significance of**<br>**difference** |
| --- | --- | --- | --- |
| EQ-5D-3L index score | 0.45 (0.34)<br>0.46 (0.34- 0.75) | 0.72 (0.22)<br>0.75 (0.58- 0.81) | P<0.001 |
| EQ-5D-3L VAS score | 49.16 (15.90)<br>50.00 (40.00- 60.00) | 57.77 (18.02)<br>60.00 (40.00- 70.00) | P<0.001 |
| SF-36 physical summary<br>score | 33.01 (12.96)<br>33.75 (24.53- 40.00) | 45.03 (15.80)<br>41.25 (32.50- 57.50) | P<0.001 |
| SF-36 Mental summary<br>score | 37.79 (10.63)<br>38.29 (32.86- 41.33) | 46.85 (13.54)<br>41.13 (37.78- 53.22) | P<0.001 |

The internal consistency reflected by the Cronbach alpha was 0.834. It is higher than all the alpha values obtained when each question is removed (Table 7).

**Table 7:** Internal consistency of the EQ-5D-3L.

| **Instance** | **Cronbach alpha** |
| --- | --- |
| When all items are present | 0.834 |
| When 1<sup>st</sup> question is removed | 0.782 |
| When 2<sup>nd</sup> question is removed | 0.815 |
| When 3<sup>rd</sup> question is removed | 0.784 |
| When 4<sup>th</sup> question is removed | 0.788 |
| When 5<sup>th</sup> question is removed | 0.832 |

## Discussion

This is the first study documenting the validity of the EQ-5D-3L among pre-dialysis patients with CKD. All the six a-priori hypotheses were found to be true in relation to this generic QOL tool. Hence this study pave the way towards opportunities of using EQ-5D-3L on patients with CKD, which is becoming a rising burden in this lower-middle income setting.

Higher proportion (i.e. more than 75%) of participants has responded as having some-or-severe problems in relation to the pain and mood domains. This is compatible with the other documented literature depicting that mental well-being of CKD affected participants must be given more emphasis(10). Additionally, it is in par with the findings of the present study with 75% being psychologically distressed and nearly 65% being depressed.

All the individual summary scores had means and medians less than 50. The figures are in general lower than the post-myocardial infarction and post-stroke QOL measurements in Sri Lanka (4,20). Comparatively the VAS values of the EQ-5D-3L too has remained around the value of 50, further proving the negative impact, CKD imposes on the lifestyles of the affected patients.

Swank and Mullen (2017) state that in interpreting convergent validity by this method, must consider three factors; significance, direction and the effect size(28). The Spearman correlation coefficients showed a significant association between the SF-36 summary scores versus EQ-5D-3L index and VAS scores in the positive direction. This shows that the parameters of EQ-5D-3L have measured the relevant constructs in the similar manner as that of SF-36 proving its construct validity. However, the effect sizes of the associations were approximately in the range of 0.2 to 0.4. This range can be classified as “acceptable and of medium strength”(28,30). The relatively weak strengths of associations are acceptable due to the high number of other factors affecting the variability of these outcome parameters. Furthermore, due to the complexities of constructs, the interpretation of findings in this regard is recommended to be not exactly similar to interpreting other bivariate correlations (31).

The index values showed relatively stronger associations compared to VAS figures. This may be explained by the fact that the index scores include more domains whereas the VAS score is a single general estimation of the participant. Since the SF-36 summary scores too include multiple domains in them, they can be assumed to be more strongly correlated with the EQ-5D-3L index score.

Both the depressed and psychologically distressed groups had relatively lower EQ-5D-3L scores compared to their counterparts. This proves that the EQ-5D-3L is valid in differentiating two groups which are known or assumed to be different in relation to QOL. The depressed and distressed people would perceive the position of their life at a lower level than those who are not. Hence the QOL of depressed and distressed groups can be assumed to be lower. In the present study EQ-5D-3L has been able to detect this difference. The SF-36 values too were mentioned to prove that actually there was a difference of QOL between the two groups compared.

The highest internal consistency was seen when all the items are present in the tool as reflected by the Cronbach alpha values. This proves that even when all the five questions are included in the tool, homogeneity of the responses is preserved. The alpha value being more than 0.7, reflects a satisfactory internal consistency as categorized conventionally (32).

There were several limitations of the study. Firstly the reliability of the EQ-5D-3L was only tested using the internal consistency in the study. To measure the reliability by test-retest method, a clinically stable period is needed (33,34). If inter-rater methods are used, a minimum time period which would eliminate the answers given to the previous responder is needed in between the data collections (35). Since EQ-5D-3L captures the QOL at the “time of completion” (22), the administration of other methods becomes debatable. On the other hand internal-consistency, though not being the sole representation, could be assumes to reflect the reliability characteristics of tools (36). However, to prevent any over-generalizability of the findings, in the title of the current study, the word “internal consistency” was used instead of the word “reliability”.

Secondly the QOL scores were not adjusted to the morbidities those were present among the participants. Majority of the participants were suffering from different comorbid conditions (nearly 71%) with different severities. Hence it was not feasible to adjust for these. However as shown in literature, concomitant non-communicable medical conditions are common among the patients with CKD (6,7) and the sample of this study too represent more or less similar characteristics.

## Conclusions

EQ-5D-3L is a valid generic QOL tool with satisfactory internal consistency to be used among CKD patients in the pre-dialysis stage

## Acknowledgement

Authors acknowledge Asanga Ranasinghe, Priyantha Kumara and Anura Ranasinghe for the support rendered during the study.

## Supporting information

S1 Appendix: Data used in the analysis

